# HOTAIR ancient sequence suggests regulatory roles both in cis and trans

**DOI:** 10.1101/250621

**Authors:** Chirag Nepal, Yavor Hadzheiv, Sachin Pundhir, Piotr Mydel, Boris Lenhard, Ferenc Müeller, Jesper B Andersen

**Author notes:** **Correspondence to:** Chirag Nepal, Biotech Research and Innovation Centre, University of Copenhagen, Ole Maaløes Vej 5, DK-2200 Copenhagen N, Denmark.

## Abstract

*HOTAIR* is a long noncoding RNA transcribed between *HOXC11* and *HOXC12* in mammals. The proposed function(s) of *HOTAIR* lacks consensus as to whether it regulates HoxD cluster genes in *trans* or HoxC cluster genes in *cis*. We have identified a 32-nucleotide long conserved noncoding element (CNE) as *HOTAIR* ancient sequence which has a paralogous copy embedded in *HOXD11* noncoding transcript. All vertebrates except teleosts have two copies of CNE and the paralogous CNEs exhibit sequence complementarity in the transcribed orientation. Moreover, paralogous CNEs underwent compensatory mutations suggesting they co-evolved and might hybridize. In both human and mouse, *HOTAIR* CNE exhibits characteristic features of a poised enhancer in *HOTAIR*-unexpressed stem cells and of an active enhancer in *HOTAIR*-expressed cells. Tight correlation between the transcriptional activity of the CNE and *HOTAIR* promoter suggests *HOTAIR* transcription is crucial for enhancer activity. In *HOTAIR-*expressed cells, *HOTAIR* expression is positively correlated with *HOXC11* in *cis* and negatively correlated with *HOXD11* in *trans*, suggesting a dual modality of *HOTAIR* ancient sequence.

## INTRODUCTION

Mammalian genomes are pervasively transcribed, giving rise to thousands of long noncoding RNAs (lncRNAs) (1,2). Only a handful of lncRNAs have well-characterized functions, which are attained through diverse mechanisms (chromatin regulation, alternative splicing, gene silencing, *cis*-regulation, *trans*-regulation) (3,4). Among these, *HOTAIR* is a prototypic model for lncRNAs regulating chromatin modifications. *HOTAIR* is an intergenic (between *HOXC11* and *HOXC12*) lncRNA proposed to regulate HOXD cluster genes (i.e., *HOXD8*, *HOXD9*, *HOXD10* and *HOXD11*) in *trans* by recruiting Polycomb Repressive Complex 2 (PRC2) (5). The proposed model has been questioned as PRC2 binding is promiscuous (6) and PRC2 is dispensable for *HOTAIR*-mediated transcriptional repression (7). Deletion of the entire mouse Hoxc cluster (including *Hotair*) showed little effect on gene expression and H3K27me3 levels at Hoxd genes (8). Specific deletion of *Hotair* produced a phenotype of homeotic transformation, skeletal malformation, global decrease in H3K27me3 levels and upregulation of posterior HoxD genes (i.e., *Hoxd10*, *Hoxd11* and *Hoxd13*) (9). These observations are now challenged as specific knockouts of *Hotair* locus *in vivo* show neither homeotic transformation nor upregulation of HoxD genes but, instead, a significant change in Hoxc (especially *Hoxc11* and *Hoxc12*) cluster genes, which argue in favor of a DNA-dependent effect of the *Hotair* deletion (10). The observed different regulatory mechanisms (*cis* versus *trans*) might be due to different tissues and developmental stages from different genetic backgrounds (11) and hence there is no consensus unifying model of *HOTAIR* regulation (12).

Some lncRNAs have enhancer-like functions (13), although it is unclear whether lncRNAs function through lncRNA transcripts or through the underlying genomic DNA with *cis*regulatory functions. Genomic deletion of lncRNA also removes *cis*-regulatory DNA elements, thus confounding whether the observed phenotype is due to the act of transcription of a lncRNA transcript or underlying genomic DNA (14). A systematic analysis involving transcription blockage and perturbation of the *Lockd* lncRNA sequence showed that it regulates *Cdkn1b* transcription through an enhancer element, while the lncRNA transcript is dispensable for *Cdkn1b* expression (15). Deletion of 12 genomic loci encoding lncRNAs revealed five loci where the deletion showed effects on the general process of transcription and enhancer-like activity but no requirement for the lncRNA products themselves (16). Identification of accurate 5’ start sites of lncRNAs revealed that a large majority of lncRNA have enhancer-like features (1). Others like *Haunt* lncRNA have dual roles where *Haunt* DNA encodes potential enhancers to activate HoxA genes and *Haunt* RNA prevents aberrant HoxA expression (17).

Here, we set to address whether *HOTAIR* regulates HOXC cluster genes in *cis* (10) or HOXD cluster genes in *trans* (5,9). HOXC and HOXD clusters are duplicated from the ancestral HOXC/D cluster, by the second round of whole genome duplication (WGD), thus we asked whether *HOTAIR* and HOXD cluster have retained some sequence from ancestral HOXC/D cluster. We have identified a 32-nucleotide conserved noncoding element (CNE) as the *HOTAIR* ancestral sequence that is present across all jawed vertebrates (except teleost) and has a paralogous copy embedded within *HOXD11* noncoding transcript. Paralogous CNEs underwent compensatory mutations and exhibit sequence complementarity. *HOTAIR* CNE represents an active or poised enhancer in different cellular context. *HOTAIR* expression is positively correlated with *HOXC11* and negatively correlated with *HOXD11*, suggesting dual modality of *HOTAIR* CNE.

## RESULTS

### Identification of *HOTAIR* ancient sequence and its paralog in HoxD cluster

Hox clusters have multiple conserved noncoding elements (CNEs) that are highly conserved from human to fish (18,19); thus, we asked whether any region of *HOTAIR* (which resides on HOXC cluster) is conserved across vertebrates. We used human and zebrafish CNEs from ANCORA (19) (see Materials and Methods) and identified a 32-nucleotide long CNE that overlapped with *HOTAIR* intronic region (Fig. 1A), which is eight nucleotides away from the splice site and overlaps a CpG island (CGI). Identification of the homolog of CNE in zebrafish is intriguing because *HOTAIR* was proposed to have a *de novo* origin in marsupials (20) and a homolog(s) of *HOTAIR* RNA have not been reported in zebrafish (21-23). Thus, we mapped the orthogonal position of the CNE in zebrafish and identified its location between *hoxd11a* and *hoxd12a* (Fig. 1B), but not in the expected hoxc clusters (Fig. 1A).

**Figure 1.**
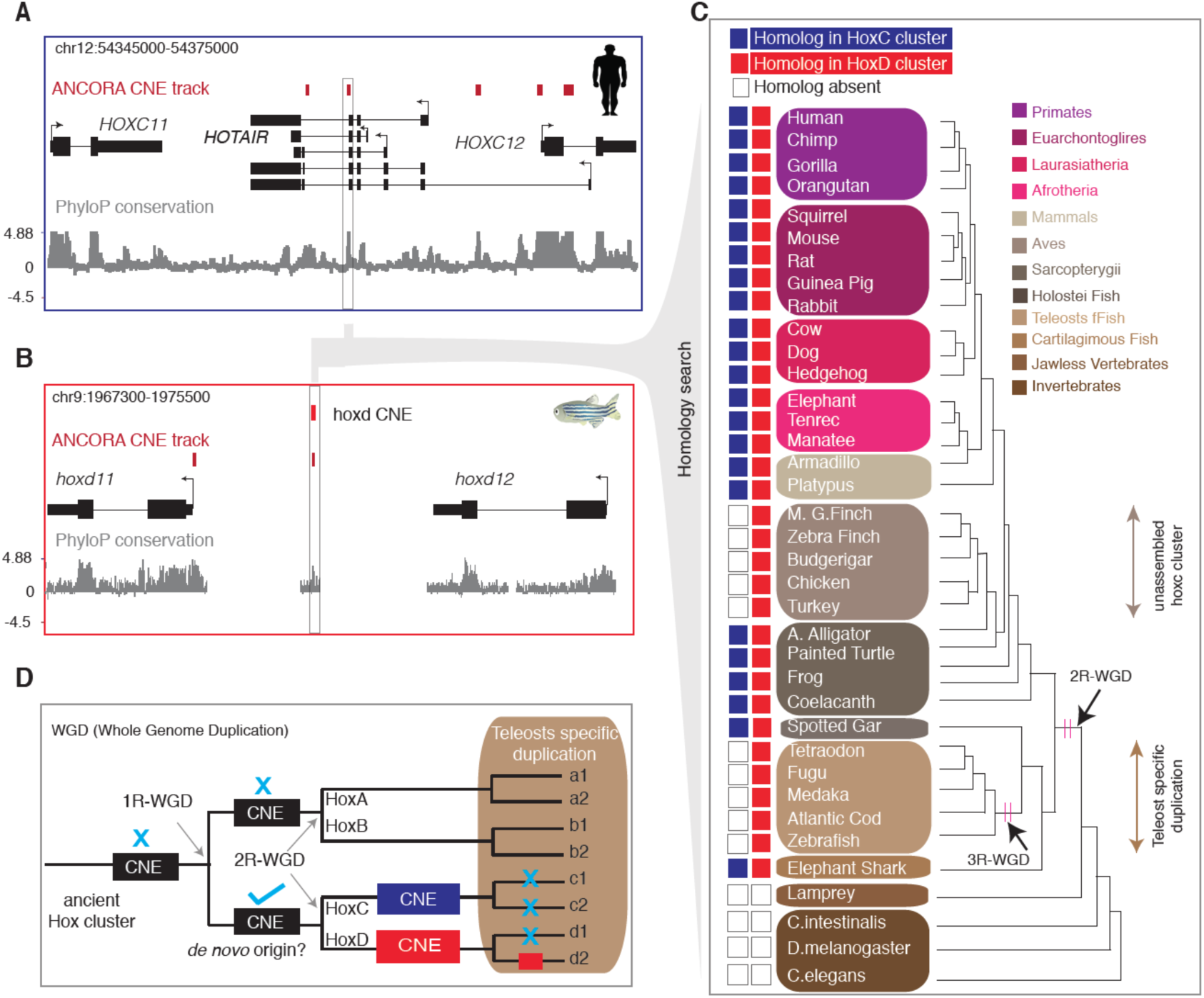
Identification of the *HOTAIR* conserved noncoding element (CNE) and its homolog in HOXD cluster across vertebrates. (**A**) A genome browser view around *HOTAIR* locus showing CNE from ANCORA browser and UCSC PhyloP conservation track. The CNE highlighted in a rectangular box is located eight nucleotides away from the splice site. (**B**) Orthologs of *HOTAIR* CNE mapped to the zebrafish hoxd (between *hoxd11* and *hoxd12*) cluster. (**C**) Homology search of the CNE across 37 species identified homologous CNEs in only HoxC and HoxD clusters. Homologs of the CNE are undetected in jawless vertebrate and invertebrates. Homologs in HoxC and HoxD clusters are represented by blue and red, respectively. Empty boxes indicate absence of homologs. (**D**) Schematic representation for the proposed model on the origin of the CNE. The CNE might have *de novo* origin in ancestral HoxC/D cluster after the first round of whole genome duplication (1R-WGD), where the second round of whole genome duplication (2R-WGD) resulted two copies of the CNE at HoxC and HoxD clusters. Additional round of teleosts specific duplication might have resulted in loss of CNE from both HoxC clusters and from one of the HoxD cluster.

As this CNE is conserved across all vertebrates (Supplementary Fig. S1A-B), we investigated whether the CNE is located in the HoxC (the capitalization “HoxC” is used to represent the HoxC cluster across multiple species) or HoxD cluster. We systematically mapped the CNE sequences across 37 organisms (34 vertebrates and 3 invertebrates) (Supplementary Table S1) and identified a homologous CNE in HoxD and HoxC clusters common to all jawed vertebrates except in teleosts and birds, but absent in lamprey (jawless vertebrate) and invertebrates (Fig. 1C). The homologous CNEs mapped between *HoxD11* and *HoxD12* in HoxD cluster and between *HoxC11* and *HoxC12* in HoxC cluster (Fig. 1C). The absence of HoxC CNE in birds is likely due to unassembled HoxC cluster (Supplementary Table S2). In contrast, teleosts have well-annotated *HoxC11* and *HoxC12* genes in the same cluster (Supplementary Table S2) but underwent an additional round of teleost-specific whole genome duplication (WGD) that might have resulted in lineage-specific loss. Thus, we conclude that all jawed vertebrates except teleosts have a homolog(s) of *HOTAIR* CNE in the HoxC and HoxD clusters.

Two rounds of WGD at the root of vertebrates have resulted in four Hox clusters (24), where the second round of WGD resulted HoxC and HoxD clusters from the ancestral HoxC/D cluster (25). Two copies of the CNE in the basal group of jawed vertebrates, such as elephant shark (cartilaginous fish) and spotted gar (basal ray-finned fish; sister group of teleosts) (Fig. 1C), suggest that the CNE was already present in the ancestral HoxC/D cluster. Therefore, it is likely that the ancestral CNE might have had a *de novo* origin in jawed vertebrates after the first WGD (Fig. 1D). However, it is also possible that the CNE was present before the first WGD and then disappeared from ancestral HoxA/B cluster. To understand how CNE flanking sequences evolved after WGD, we aligned flanking sequences within species and observed limited homology (for example, in human and elephant shark; Supplementary Fig. S2A) suggesting only CNEs were under selection. Then we separately aligned HoxC and HoxD CNE flanking sequences and observed independent conservation along each cluster (Supplementary Fig. S2B) and, in turn, identified a relatively long stretch of about 175 nucleotides of *HOTAIR* sequence that is conserved across vertebrates (except teleosts).

### Paralogous HoxD CNE is transcribed from *HoxD11* noncoding transcript and exhibit sequence complementarity with *HOTAIR* CNE

Some CNEs are transcribed from noncoding transcripts (26); thus, we asked if human HOXD CNE is transcribed from coding/noncoding transcript. We intersected the genomic coordinates of the CNE with Ensembl genes and observed HOXD CNE is transcribed from an alternative noncoding isoform of *HOXD11* (referred to as *HOXD11* noncoding transcript) (Fig. 2A), which is a reported target gene that is silenced by *HOTAIR* in both human (5) and mouse (9). As both CNEs are embedded within their respective transcripts, we asked whether these CNEs exhibit sequence complementarity, similar to a miRNA seed and target sites. We aligned paralogous CNEs in the orientation of their respective transcripts and observed sequence complementarity (Fig. 2A), suggesting paralogous CNEs might hybridize.

**Figure 2.**
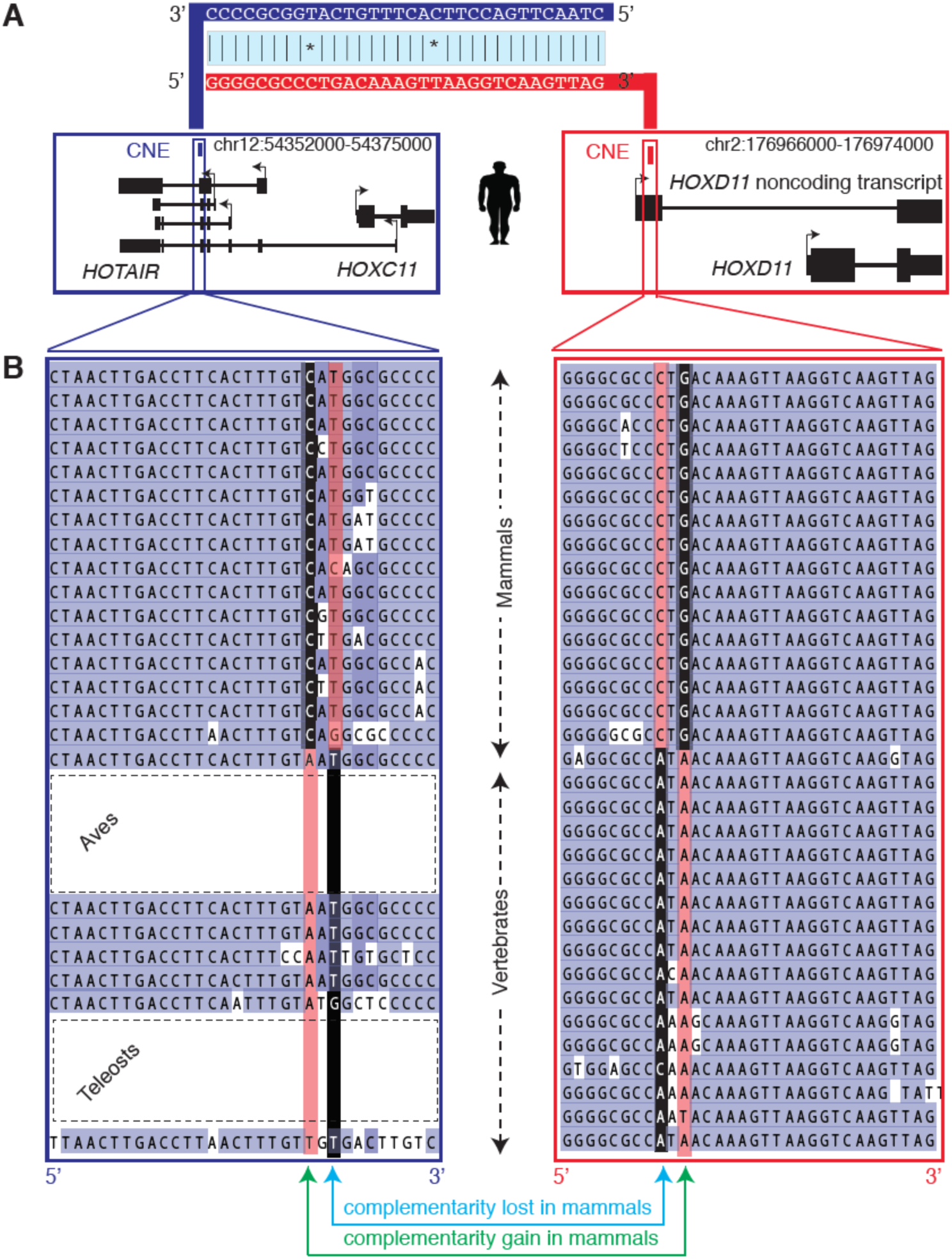
Sequence complementarity of the *HOTAIR* CNE and the paralogous HoxD CNE. (**A**) *HOTAIR* CNE and HOXD CNE are represented by rectangular blue and red bars, respectively. *HOTAIR* CNE and HOXD CNE are zoomed and show the genomic DNA sequences. *HOTAIR* is transcribed from negative strand and the CNE sequence is reverse transcribed to show in 5’ to 3’ orientation. Paralogous CNEs are complementary in the orientation of transcripts suggesting they might hybridize if co-expressed in the same cells. (**B**) Alignment of the *HOTAIR* CNE and HoxD CNE in transcribed orientation reveal sequence complementarity is conserved across vertebrates. Genetic substitution within paralogous CNEs occurred at specific positions that co-evolved in two successive waves in vertebrates and mammals resulting in gain of complementarity at one position and loss of complementarity at other position.

To confirm whether the observed sequence complementarity in human is conserved in other vertebrates, we analyzed Ensembl-annotated genes and RNA-seq transcripts (22,27) across a subset of species and identified the *HOTAIR* and *HoxD11* noncoding transcripts in mouse, ferrets, dog and coelacanth (Supplementary Fig. S3A-B). The *HoxD11* noncoding transcript was detected in chicken but undetected in teleosts (zebrafish and tetraodon; Supplementary Fig. S3A-B). For other species, we inferred the putative orientation of the missing transcript (see Materials and Methods), as illustrated for chimp and painted turtle (Supplementary Fig. S3C-D). We aligned CNEs in the 5’ – > 3’ orientation and observed two interesting patterns. First, the sequence complementarity observed in human is conserved across vertebrates (Fig. 2B). This suggests that sequence complementarity between paralogous CNEs is an ancient feature that has been under selection pressure for more than 300 million years. This raises an important question as to whether the key function of these transcripts is to provide transcription of the CNE. Second, both CNEs underwent substitutions at specific positions that evolved in two separate waves in vertebrates and mammals (Fig. 2B). One of the substitutions resulted in gain of complementarity in mammals and the other substitution resulted in loss of complementarity in mammals (Fig. 2B). Collectively, the co-evolution of CNEs and retention of sequence complementarity in the transcribed orientation suggest that paralogous CNEs might interact and be co-regulated.

### *HOTAIR* CNE reflects a poised enhancer in *HOTAIR-* unexpressed stem cells

CNEs are putative regulatory elements (28) where many of them function as enhancers (29); we thus asked whether CNEs have open chromatin state as in active regulatory elements (30,31). We selected 29 cell lines (Supplementary Table S3; see Materials and Methods) from Roadmap Epigenome (30) and classified them as *HOTAIR*-expressed (N=10) and *HOTAIR*-unexpressed (N=19) groups based on RNA-seq expression levels (Supplementary Fig. S4A). Among *HOTAIR*-unexpressed cells, H1-hESC cell line is unique as the CNE is enriched for both repressive (H3K27me3) and open chromatin (as indicated by DNase hypersensitive sites (DHSs)) marks (Supplementary Fig. S4B), suggesting it to be in the poised state. We observed enriched DHS signals around the CNE in *HOTAIR*-expressed cells and *HOTAIR*-unexpressed stem cells, but not in *HOTAIR*-unexpressed differentiated cells (Fig. 3A). The paralogous HOXD CNE also has a similar chromatin state in corresponding cell lines (Supplementary Fig. S4C). Similar analysis on mouse cell lines (32) (see Materials and Methods) revealed that open chromatin state around the CNE is conserved in *Hotair*-expressed cells and *Hotair*-unexpressed stem cells (Fig. 3A; Supplementary Table S3). The chromatin state of *Hotair* CNE gradually becomes open during reprogramming of mouse embryonic fibroblasts (MEF) to induced pluripotent stem cells (iPSC) (33) (Supplementary Fig. S4D). We conclude that the chromatin state of CNE is open in *HOTAIR*-unexpressed stem cells and *HOTAIR*-expressed cells in both human and mouse.

**Figure 3.**
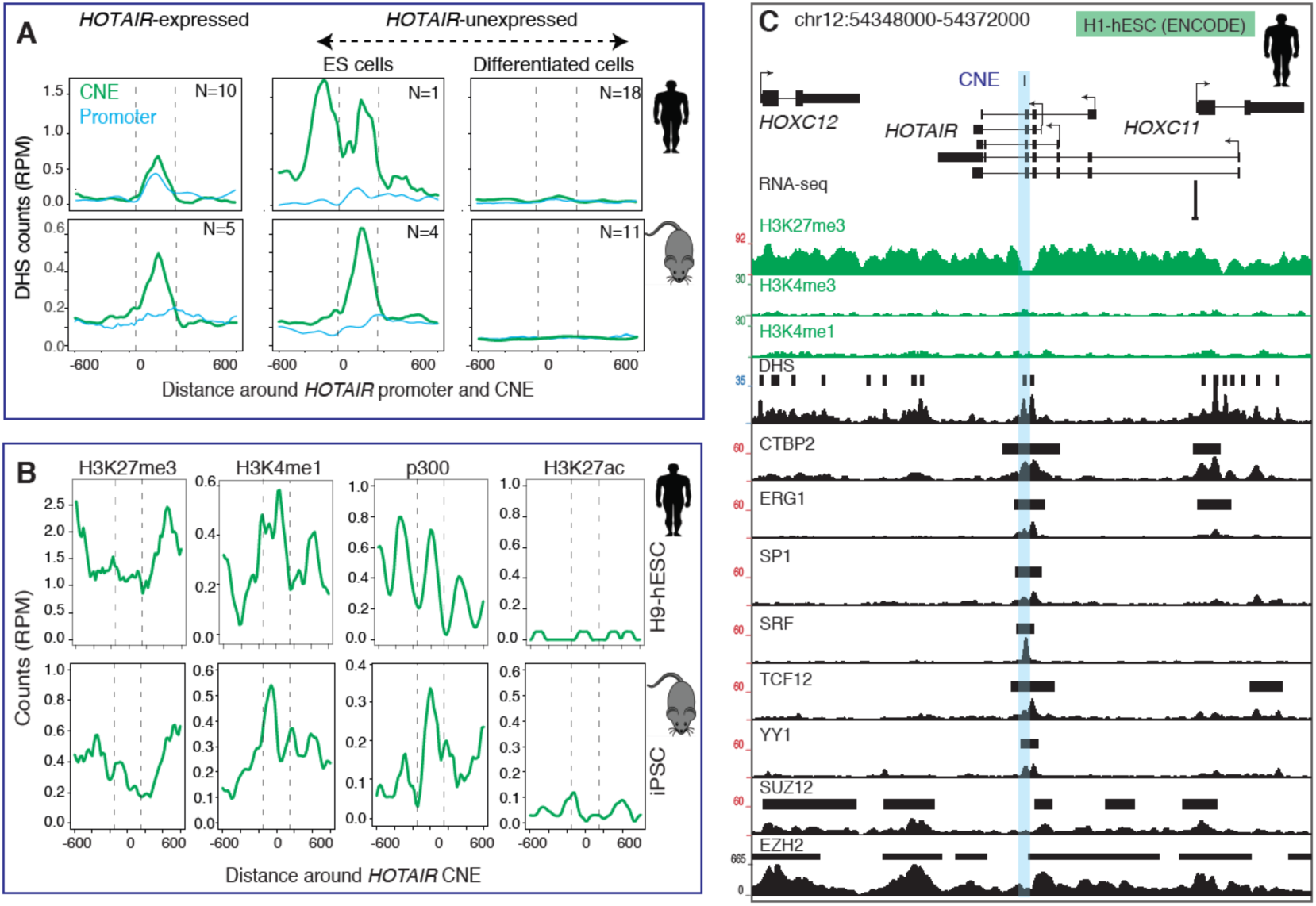
The *HOTAIR* CNE represents a poised enhancer in *HOTAIR*-unexpressed stem cells in both human and mouse. (**A**) The *HOTAIR* CNE chromatin is open in *HOTAIR*-expressed and *HOTAIR*-unexpressed stem cells in human and mouse. Y-axis is normalized DNase I hypersensitive sites (DHS) coverage in reads per million (RPM). “N” denotes the number of cell lines. (**B**) The *HOTAIR* CNE is marked by p300, H3K4me1 and bimodal H3K27me3 peaks in H9-hESC (human) and iPSC (mouse) cell line that collectively define a poised enhancer. Y-axis is normalized coverage in reads per million (RPM). (**C**) A genome browser view around the *HOTAIR* locus with transcription factors, DHS, histone modifications and RNA-seq tracks from H1-hESC cell line. *HOTAIR* is unexpressed (as shown by RNA-seq track) and marked by broad H3K27me3 peak. The CNE is marked by depleted H3K27me3 marks, reflecting a nucleosome-depleted region which is supported by enriched DHS peaks and binding of multiple transcription factors.

A subset of embryonic stem cell-specific enhancers are poised, characterized by presence of both activating and repressing marks (34); thus, we asked whether CNE reflects a poised enhancer in *HOTAIR*-unexpressed stem cells. We observed that H3K4me1, H3K27me3 and p300 signals are enriched at the CNE (Fig. 3B) in human H9-hESC (34) and mouse iPSC (33), supporting that *HOTAIR CNE* is a poised enhancer. However, the HOXD CNE lacks p300 and bimodal H3K27me3 peaks (Supplementary Fig. S4E). Poised enhancers have bimodal H3K27me3-modified nucleosomes (34), which is observed in *HOTAIR* CNE (H1-hESC and H9-hESC) (Fig. 3B; Supplementary Fig. S4F) but not in HOXD CNE.

We next analyzed all ENCODE transcription factors (TFs) (35) and observed CTBP2, YY1, SP1 and CHD1 are specifically enriched on the CNE exclusively in H1-hESC (Fig. 3C; Supplementary Table S4). YY1 has a known dual function in repressing or activating transcription depending upon the cofactors it recruits (36). The binding of YY1 on regulatory elements and their associated RNA species has been shown to play a role at enhancers (37). PRC2 components (SUZ12 and EZH2) are marked throughout the locus (Fig. 4C) that are not specific to stem cells (Supplementary Table S4). A recent study on mouse embryonic stem cells using promoter capture Hi-C data showed extensive promoter contacts for HoxC and HoxD clusters (38). Poised enhancers establish physical interactions with their target genes in ESCs in a PRC2-dependent manner (39), whether *HOTAIR* CNE might have a similar role in stem cells remains speculative. We conclude that in both human and mouse, the *HOTAIR* CNE possess characteristics features of a poised enhancer in stem cells.

### *HOTAIR* CNE reflects an active enhancer in *HOTAIR-* expressed cells

Bidirectional transcription of enhancer RNAs (eRNAs) is a hallmark of active enhancers (40). We therefore analyzed the FANTOM5 (41) CAGE signals at the CNE and observed bidirectional transcription flanking the CNE (Fig. 4A; dashed rectangular box), supporting its role as an active enhancer. The CNE and the promoter region of *HOTAIR* primary and alternative transcripts are co-expressed (Fig. 4B; Supplementary Table S5), as exemplified during differentiation of myoblast to myotube (Supplementary Fig. S5A). The expression of *HOTAIR* promoter and CNE are positively correlated (Fig. 4C), which also holds true for two other alternative promoters except for the distal promoter (labelled as dp1) (Supplementary Fig. S5B). Negative correlation of distal promoter is mostly due to large number of samples where only distal promoter is expressed (Fig. 4B). The genomic region between bidirectional CAGE tags around the CNE is about the length of one nucleosome (Fig. 4D) and is conserved across vertebrates (Supplementary Fig. S2B). DHS, H3K4me1 and H3K27ac peaks are also enriched around the CNE (Fig. 4A) in myoblast and myotube (30), thus providing additional evidence as an active enhancer. The CNE is flanked by bimodal H3K4me1 peaks, a characteristic feature of active enhancers (Fig. 4E). Due to lack of appropriate cell lines (embryonic hindlimbs, genital tubercle and piece of trunk corresponding to the sacro-caudal region) where *Hotair* is expressed (8-10), we did not detect CAGE tags in either *Hotair* CNE or promoter across mouse FANTOM5 samples (Supplementary Fig. S5C). However, we observed that H3K4me1 and H3K27ac are enriched in mouse embryonic (E10.5 days) hindlimbs (42) (Fig. 4F; Supplementary Fig. S5D).

**Figure 4.**
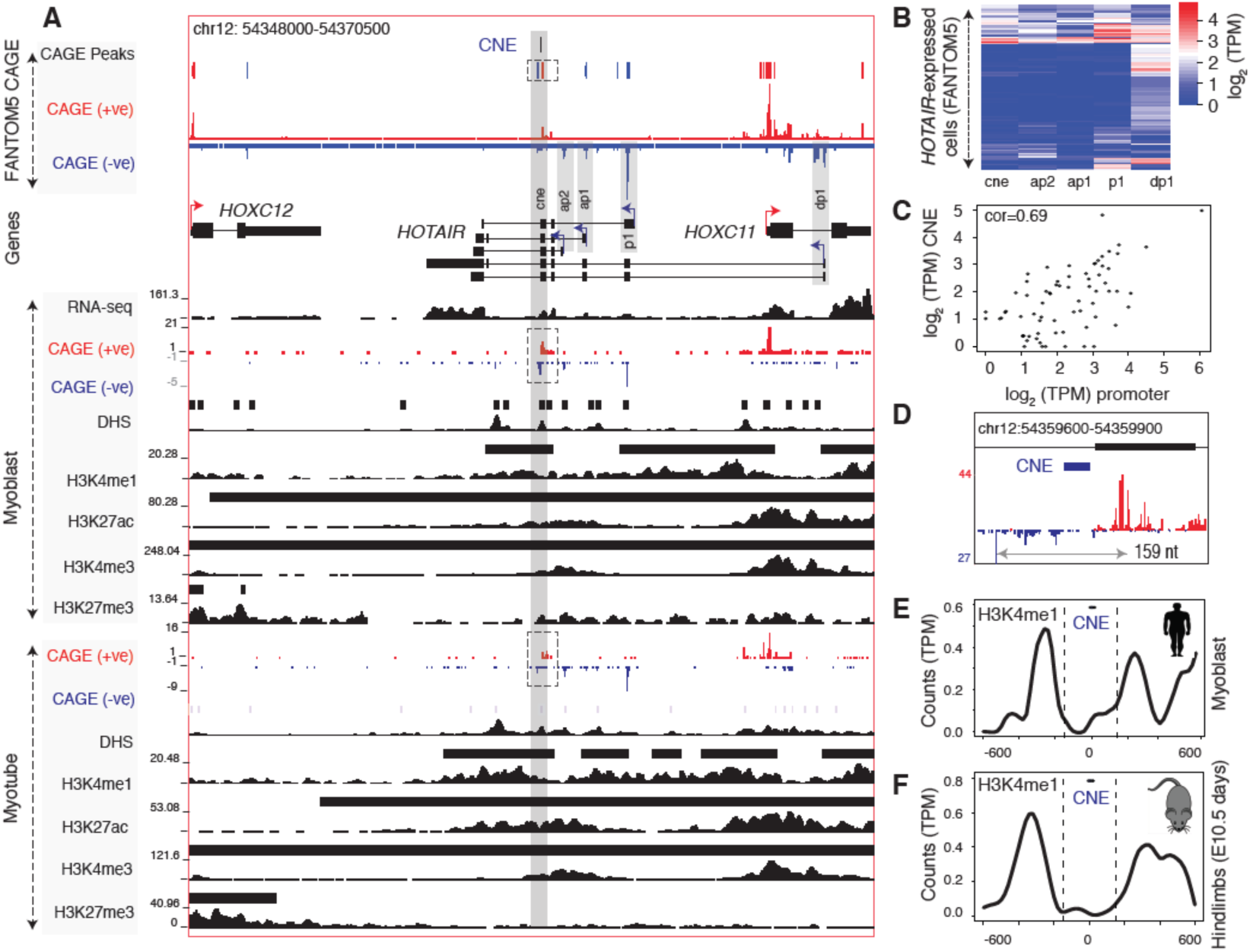
The *HOTAIR* CNE reflects an active enhancer in *HOTAIR*-expressed muscle cells. (**A**) A genome browser view with FANTOM5 CAGE tags (combined tracks) and individual tracks on myoblast and myotube along with RNA-seq, H3K4me/me3, H3K27ac/me3, and DNase hypersensitive site (DHS) tracks from ENCODE. Horizontal bars across each histone and DHS tracks are annotated peaks. Forward and reverse strand CAGE tags are represented by blue and red. Bidirectional CAGE tags flanking the *HOTAIR* CNE is shown in dashed rectangular boxes. In both myoblast and myotube, the *HOTAIR* CNE is flanked by bidirectional CAGE tags and overlaps H3K4me1 and H3K27ac peaks. (**B**) Expression levels of the *HOTAIR* CNE and alternative promoters across FANTOM5 samples. (**C**) Expression level of the CNE is correlated with the *HOTAIR* promoter. (**D**) A zoomed view of CAGE tags (from myoblast and myotube differentiation time points) around the CNE shows bidirectional CAGE peaks that roughly represent the length of a nucleosome. (**E-F**) Bimodal H3K4me1 peaks flank the CNE in human myoblast (**e**) and embryonic 10.5days hindlimbs in mouse (**F**).

Thus, we have shown that *HOTAIR* CNE possesses characteristic features of an active enhancer in *HOTAIR*-expressed cells, in both human and mouse. Importantly, transcription of the *HOTAIR* promoter is tightly linked to enhancer activity of the CNE, suggesting transcription of the CNE is a key function of the *HOTAIR* transcript. This notion is further supported by the evolution of CNE paralogs that reveal transcribed paralogous CNEs have retained sequence complementarity and thus have been actively selected during evolution (Fig. 2; Supplementary Fig. S3C-D). This observation suggests that transcription directly on the CNE might be a key contributor to the selection acting on the CNE.

### *HOTAIR* CNE expression is positively correlated with *HOXC11* and negatively correlated with *HOXD11*

The lack of consensus on whether *HOTAIR* regulates the HoxD cluster posterior (*HoxD11*, *HoxD12*) genes *in trans* (5,9) or HoxC cluster (*Hoxc10*, *Hoxc11*) genes *in cis* (10) prompted us to ask how the expression levels of *HOTAIR* CNE is correlated with reported target genes in large samples of *HOTAIR*-expressed cells. We reasoned that *HOTAIR* CNE reflects an active enhancer and could potentially explain *cis* regulation, while the sequence complementarity between *HOTAIR* CNE and HOXD CNE (embedded within *HOXD11*) could potentially regulate HOXD cluster genes in *trans*. To this end, we analyzed 694 cell types from FANTOM5 (41) and plotted the expression levels of genes across four HOX clusters (Supplementary Fig. S6A). We next selected only the *HOTAIR*-expressed cells and observed positive correlation with HOXC posterior genes and a trend towards negative correlation with HOXD posterior genes (Fig. 5A). It is striking to note that among HOXC and HOXD cluster posterior genes, *HOXC11* expression has the best positive correlation (R=0.64; p-value: 2.2E-13) in the HOXC cluster and *HOXD11* coding (R=-0.25; p-value: 0.009) and noncoding transcript (R=-0.23; p-value: 0.017) has the best negative correlation, both of which are target genes regulated by *HOTAIR* in human (5) and mouse (9,10). The observed negative correlation on HOXD cluster genes will diminish if analyzed by combining *HOTAIR*-expressed and *HOTAIR*-unexpressed cells. To confirm whether the observed correlation is present in other datasets, we analyzed gene expression data of breast cancer patients from TCGA (43). We selected only those patients (N=605) where *HOTAIR* is expressed (see Materials and Methods) and observed a similar trend of positive correlation with *HOTAIR* and HOXC cluster posterior genes and a trend of weak negative correlation with *HOTAIR* and HOXD cluster posterior genes (Supplementary Fig. S6B). Similarly, *HOXC11* has the highest positive correlation (R=0.207; p-value:2.63E-07) and *HOXD11* has highest negative correlation (R=-0.107; p-value:0.008). The act of transcription by *HOTAIR,* transcribes the *HOTAIR* CNE (putative enhancer) and collectively regulates neighboring genes *in cis*. On the other hand, sequence complementarity of *HOTAIR* CNE with paralogous HOXD CNE embedded in the *HOXD11* noncoding transcript might negate the expression of *HOXD11*. Our observations suggest that *HOTAIR* CNE might simultaneously regulate *HOXC11 in cis* and *HOXD11 in trans*.

**Figure 5.**
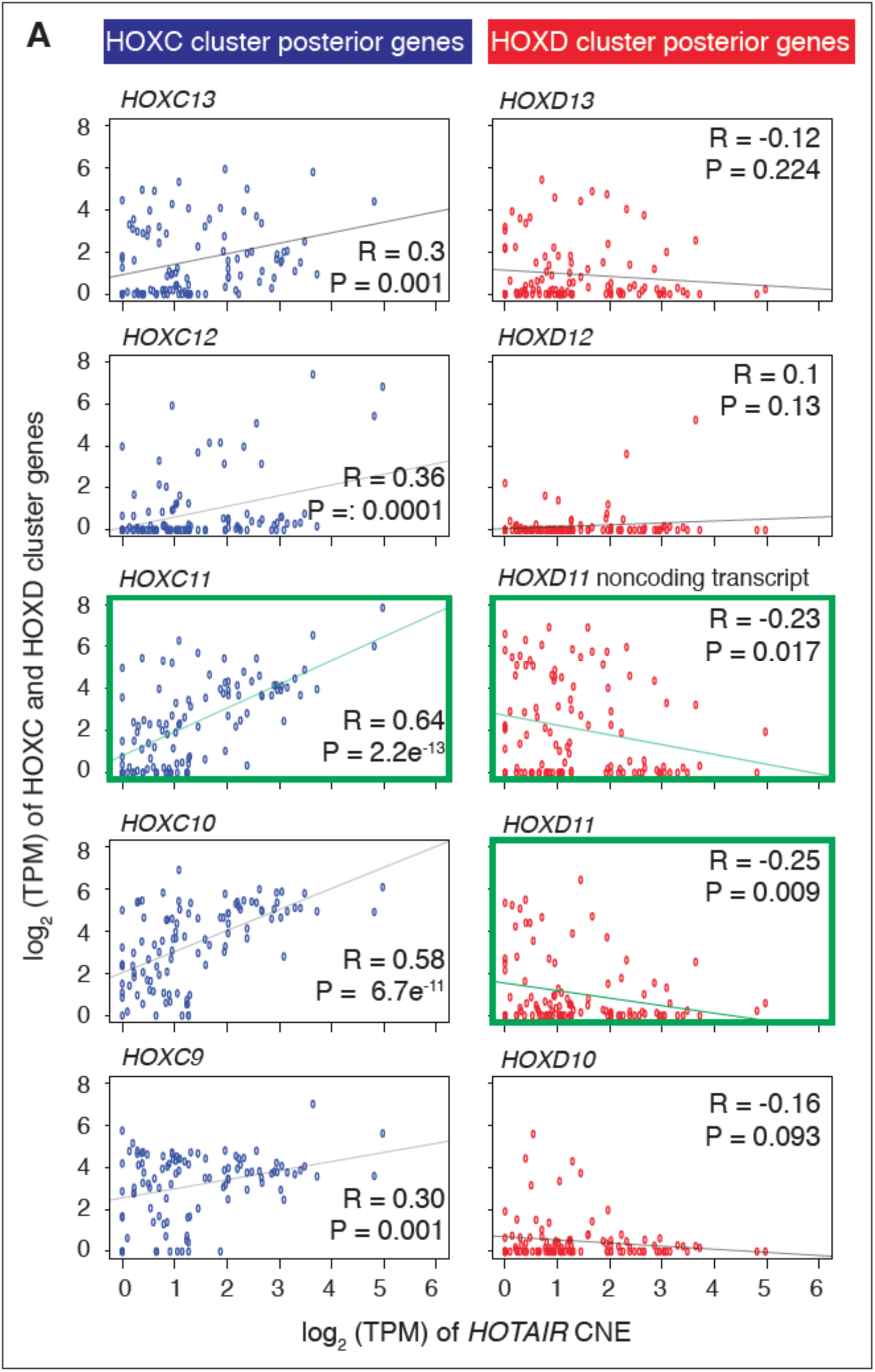
Correlation of expression levels of *HOTAIR* CNE with HOXC and HOXD clusters posterior genes in *HOTAIR*-expressed cells across FANTOM5 samples. (**A**) Expression levels of *HOTAIR* CNE with HOXC and HOXD cluster posterior genes on individual cell types. X-axis represents expression level of *HOTAIR* CNE and y-axis represents the expression levels of HOXC and HOXD cluster genes. Expression level is measured as tags per million (TPM). Expression levels of *HOTAIR* CNE and HOXC cluster genes show a trend of positive correlation where *HOXC11* (highlighted in green) has the highest positive correlation. Expression levels of *HOTAIR* CNE and HOXD cluster genes showed a trend of weak negative correlation where *HOXD11* coding and noncoding (highlighted in green) is the most negatively correlated gene.

## DISCUSSION

We have identified and characterized the *HOTAIR* CNE as the ancestral sequence, which also has a paralogous coy in the HoxD cluster across vertebrates. Lack of the CNE in teleosts HoxC cluster is unexplained given it is conserved in other fish (spotted gar and elephant shark) and is likely one of the consequences of teleosts lineage whole genome duplication. Conservation of *HOTAIR* CNE and flanking sequences even in early fish (elephant shark, spotted gar) and tetrapod (coelacanth) (Supplementary Fig. S2B) does suggest an ancestral form of *HOTAIR* across all vertebrates, but needs additional RNA-seq transcript evidence which is currently limited to only few species due to limited data. Importantly, the length of the *HOTAIR* CNE flanking sequences that is conserved across vertebrates (Supplementary Fig. S2B) roughly represents region of DHS peak and bidirectional CAGE tags.

**Figure 6.**
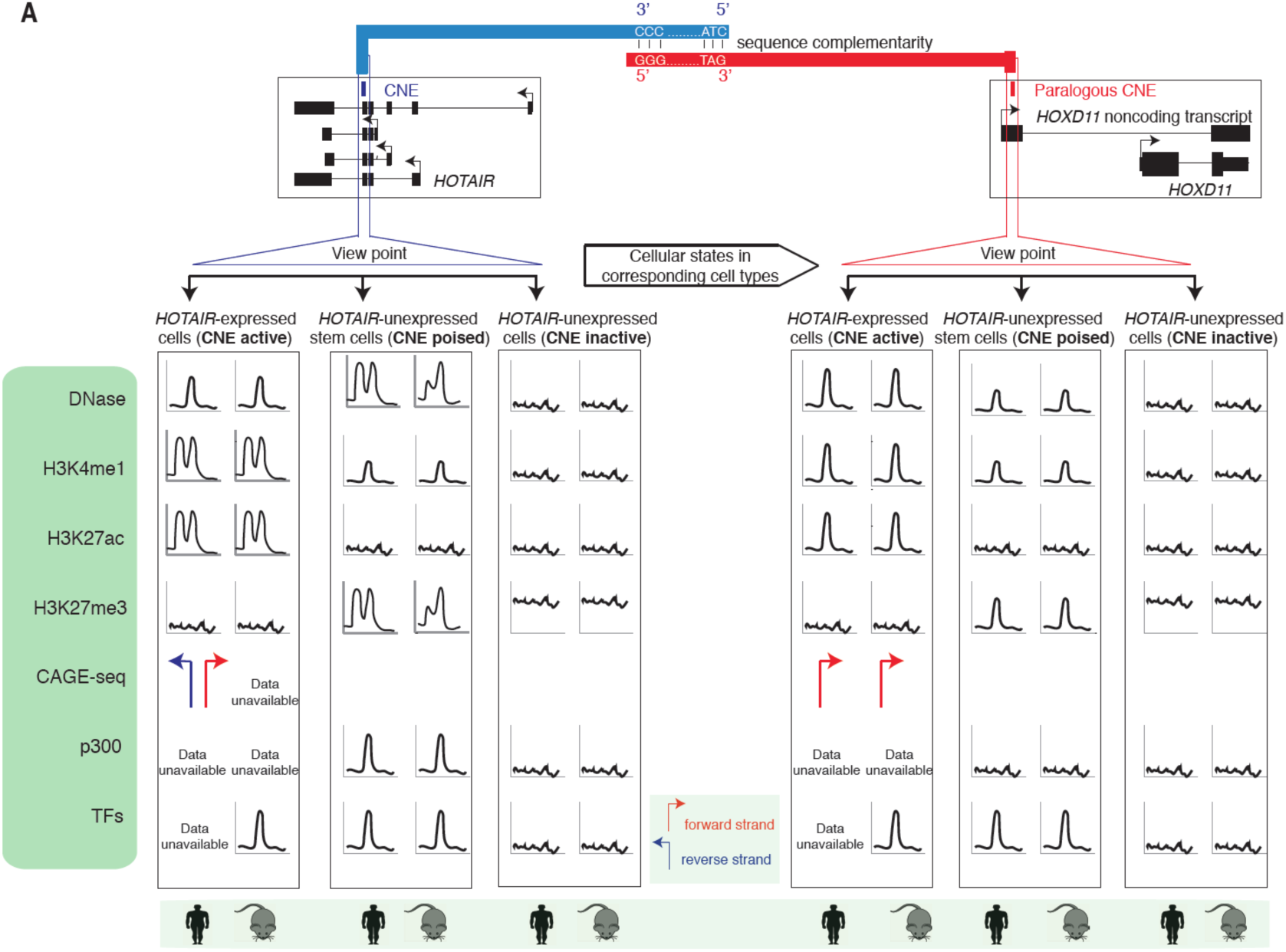
Schematic representation of various features of the *HOTAIR* CNE and its paralog in HoxD cluster in human and mouse. (**A**) The *HOTAIR* CNE has a paralog in HoxD cluster. Transcribed paralogous CNEs exhibits sequence complementarity and this complementarity is conserved across vertebrates. Based on *HOTAIR* expression, cell types are divided into *HOTAIR*-expressed and *HOTAIR*-unexpressed groups, where HOTAIR-unexpressed cells are further divided into stem cells and terminally differentiated cells based on DHS signals. Chromatin states, transcription factor bindings and CAGE signals on homologous CNEs are shown schematically.

Many of the tested CNEs act as enhancers in reporter construct (29). In fact, the homologous Hoxd CNE drives expression in a proximal posterior part of developing forelimbs (44) but does not work in its own context in zebrafish, as it requires the mouse promoter. However, deletion ofHoxd CNE in *vivo* revealed no phenotype (45), and it was speculated that the difference might be due to other phenotypes that were not detected or has a redundant copy of the sequence that have masked the effect. We have now identified a paralog of the Hoxd CNE in *Hotair* (Fig. 1). This raises an interesting question whether paralogous CNEs have redundant functions, such that deletion of one CNE might be compensated by the other CNE. Putting this into the context of deletion of *Hotair* locus (including CNE) *in vivo* (9,10), it remains unknown whether the effects of *Hotair* CNE deletion is compensated for, to a certain extent, by paralogous Hoxd CNE. Duplicated CNEs have retained only a single copy (28); thus, retention of both copies of duplicated CNE is rare and provides an interesting model to test redundancy/sub-functionalization of CNEs.

Mechanistically, *HOTAIR* was proposed to regulate HOXD cluster genes by recruiting PRC2 (5), but this model has now been questioned since PRC2 is dispensable from *HOTAIR-*mediated transcriptional repression (7). Our observation of sequence complementarity between *HOTAIR* CNE and HOXD CNE (Fig. 2; Supplementary Fig. S3A-B; Fig. 6), suggest sequence-specific binding. This notion is further supported by the observed simultaneous genetic substitutions of CNEs during evolution. Moreover, paralogous CNEs are generally coexpressed in the same cell lines (Fig. 6a), simultaneously maintaining open or closed chromatin (Fig. 6) and expression of the *HOTAIR* CNE and *HOXD11* (previously reported target gene of *HOTAIR* (5,9)) is negatively correlated which collectively provide strong arguments that these CNEs might interact.

*HOTAIR* has been the prototypic model of functional lncRNA and, thus, the presence of an embedded enhancer (characterization based on bidirectional CAGE tags, DHS and histone peaks) was unexpected (Fig. 6). Most importantly, transcriptional activity of the CNE is coupled to *HOTAIR* transcription, suggesting that a key function of the *HOTAIR* transcript is to provide active transcription for the CNE. Though the expression of the *HOTAIR* CNE is positively correlated with *HOXC11* (Fig. 5), it remains unknown whether the observed *cis* regulation of *Hotair* (10) is mediated through the CNE. However, we observed that *HOTAIR* expression is positively correlated with *HOXC11* and negatively correlated with *HOXD11* and suggest a unifying model of *HOTAIR* regulation through CNE. Our findings have opened an avenue for multiple testable hypotheses that should clarify ongoing controversies whether *HOTAIR* regulates HoxC or HoxD cluster genes *in cis* or *trans* (5,7-10) or can regulate both HoxC and HoxD cluster genes simultaneously. Our results should guide future experiments to resolve ongoing controversies. Future studies should focus on understanding how specific deletion of CNE(s) alters *HOTAIR* regulation in different cellular contexts.

## MATERIALS AND METHODS

### Genome assemblies and gene annotations

Analyses on human and mouse were done in hg19 and mm9 genome version respectively. The genome assemblies of 37 species are listed in Supplementary Table S1. Gene models were downloaded from UCSC (46). The conserved noncoding elements (CNEs) were downloaded from ANCORA (19).

### Roadmap Epigenome, ENCODE, mouse ENCODE and FANTOM5 data sets

Roadmap Epigenome data were downloaded from NIH Roadmap Epigenome browser (30). Annotated chromatin states were downloaded from 127 cell lines. Histone modifications (H3K4me1/3, H3K27ac/me3) and DNase I hypersensitive sites (DHSs) data were downloaded as mapped (tagAlign format) files. RNA-seq data for 57 cell lines were downloaded in the computed gene expression (RPKM) matrix (30). Only 29 cell lines that have all four (H3K4me1/3, H3K27ac/me3) histone modifications, DHS and RNA-seq were used for downstream analysis.

Histone modification and RNA-seq data from ENCODE were downloaded as mapped BAM files. ENCODE transcription factor ChIP-seq were downloaded as annotated peaks (35). Histone modifications (H3K4me1/3, H3K27ac/me3), DHS and RNA-seq data from mouse ENCODE were downloaded as mapped BAM files (32). Samples with replicates were merged into a single file. A threshold of 0.5 RPKM (reads per kilobase per million) was used as the cutoff expression to determine whether *HOTAIR* is expressed or not in the given RNA-seq samples.

Human and mouse CAGE-seq data were downloaded from FANTOM5 (41,47). Replicates were pooled into single file and resulting CAGE tags in each sample were quantified as tags per million (TPM) (21). CAGE tags with the highest expression level defined dominant transcription start site (TSS). CAGE based expression level was computed by summing all CAGE tags in defined the promoter region (300 bases upstream and downstream of TSS). To compare the expression correlation across four HOX clusters genes, we selected only 694 cell types only if any of HOX genes was expressed at the minimum of 5 TPM.

### Data sets used from multiple studies

RNA-seq transcripts for multiple species were used from previous studies(21,22,27). Raw data during reprogramming of mouse embryonic fibroblast to iPSC were download from GEO (GSE90894) (33). Raw data for H3K27me3, H3K4me1, H3K27ac and p300 from H9-hESC cell lines were download from GEO (GSE24447) (34). Mouse embryonic (10.5 days) hind limb data were downloaded from GEO (GSE84793) (42). Raw fastq reads were mapped using bowtie2 (48). Only unique mapping reads were considered for downstream analysis. Breast cancer patients mRNA (Illumina Human v3 microarray) data (43) were downloaded from TCGA portal. Expression levels are measured in z-scores. We filtered samples where *HOTAIR* expression ( <= 0.5) and were left with 605 patients.

### Mapping of *HOTAIR* CNE across multiple species

The *HOTAIR* CNE sequences from both human and zebrafish was used as a query sequence. We used BLAST (blastall –p blastn –d –e 0.01 –m 8)(49) to find homologous sequences against 37 species (Supplementary Table S1). Even at the permissive e-value cutoff of 0.01, only two homologous sequences were identified.

### Determining the orientation of transcripts overlapping CNEs

The *HOTAIR* transcript is annotated in multiple species (22,46), such as human, chimp, mouse, ferret and dog, and its orientation is antisense to *HoxC11* and *HoxC12* genes. Species lacking *HOTAIR* annotation, orientation of *HOTAIR* CNE was assigned antisense to annotated HoxC cluster genes. In multiple species, such as human, chimp, mouse, ferret, dog and chicken, HoxD CNE is embedded within the untranslated region of *HoxD11* noncoding transcript, which is an alternative splice variant of *HoxD11* coding gene. Thus, orientation of HoxD CNE was assigned similar to that of annotated *HoxD11* gene. Among teleosts fish, we analyzed RNA-seq transcripts in zebrafish(21,22) and tetraodon (27), and did not identify *HoxD11* noncoding transcript.

### Software and tools

Multiple alignments were generated using ClustalW (50) and Jalview (51). Sequence logos were generated using WebLogo (52). Visualization of multiple data were done by uploading bigwig tracks on UCSC genome browser and images were downloaded. Bedtools (53), bash, perl and R scripts were used for data analysis.

## ACKNOWLEDGEMENTS

We are very grateful to ENCODE, FANTOM5, Roadmap Epigenome consortia and other researchers for making data freely available. We thank Colm J. O’Rourke, Michal Lubas and Albin Sandelin for critical comments on the manuscript. C.N acknowledges Boris Lenhard, Vidar Martin Steen, Piotr Mydel and Jesper B Andersen for supporting him.

## AUTHOR’S CONTRIBUTION

C.N conceived the story, analyzed data and interpreted results. Y.H, S.P, P.M, B.L, F.M and J.B.A contributed to data analysis and critical discussions. All authors contributed to writing the manuscript.

## SUPPLEMENTARY TABLES LEGENDS

**Supplementary Table 1.** The genomic coordinates of the CNE in HoxC and HoxD cluster across species. Lack of homolog is denoted by “-”. The orientation of the *HOTAIR* CNE is antisense to the annotated *HoxC11* gene and the orientation of the transcript overlapping HoxD CNE is similar to that of annotated *HOXD11*. Number of mismatches between paralogous CNEs is shown. Evidence of Ensembl or RNA-seq annotated host transcript-encoding CNEs is denoted by “Y” and if there is no evidence, as in case of teleosts it is denoted by “N”. No evidence of transcripts due to lack of data is denoted by “NA”.

|  | Coordinates of CNE homologs |  | Count | Host Transcript Evidence |  |
| --- | --- | --- | --- | --- | --- |
|  | HoxC cluster<br>Chr:Start-End(Strand) | HoxD cluster<br>Chr:Start-End(Strand) | Mismatches | HoxC | HoxD |
| **Primates** |  |  |  |  |  |
| Human(hg19) | chr12:54359707-54359739(-) | chr2:176969157-176969189(+) | 2 | Y | Y |
| Chimp(panTro4) | chr12:35284661-35284693(+) | chr2B:180579781-180579813(+) | 2 | Y | Y |
| Gorilla(gorGor3) | chr12:52012160-52012192(-) | chr2B:64029540-64029572(+) | 3 | Y | Y |
| Orangutan(ponAbe2) | chr12:53740632-53740664(-) | chr2b:66260351-66260383(+) | 3 | Y | Y |
| **Euarchontoglires** |  |  |  |  |  |
| Squirrel(speTri2) | JH393322:11187590-11187622(-) | JH393319:5591308-5591340(+) | 3 |  |  |
| Mouse(mm10) | chr15:102947454-102947486(-) | chr2:74679612-74679644(+) | 4 | Y | Y |
| Rat(rn5) | chr7:142349588-142349620(-) | chr3:68068756-68068788(+) | 4 |  | Y |
| GuineaPig(chiLan1) | scaffold_9:45845103-45845135(+) | scaffold_3:14206941-14206973(-) | 4 |  |  |
| Rabbit(oryCun2) | chr4:37750144-37750176(-) | chr7:115664606-115664638(+) | 3 |  |  |
| **Laurasiatheria** |  |  |  |  |  |
| Cow(bosTau8) | chr5:26225443-26225475(+) | chr2:20844410-20844442(-) | 2 |  |  |
| Dog(canFam3) | chr27:1296692-1296724(+) | chr36:19912089-19912121(+) | 3 | Y | Y |
| HedgeHog(eriEur3) | JH835374:2711697-2711729(+) | JH835538:1267467-1267499(+) | 4 |  |  |
| **Afrotheria** |  |  |  |  |  |
| Elephant(loxAfr3) | scaffold_2:40399191-40399223(-) | scaffold_3:51280894-51280926(-) | 3 | NA | NA |
| Tenrec(echTel2) | JH980427:730939-730971(-) | JH980302:1668675-1668707(+) | 4 | NA | NA |
| Manatee(triMan1) | JH594633:22473672-22473704(+) | JH594643:8289255-8289287(-) | 3 | NA | NA |
| **Mammals** |  |  |  |  |  |
| Armadillo(dasNov3) | JH569064:163083-163115(+) | JH575655:3643342-3643374(+) | 0 |  |  |
| Platypus(ornAna1) | Contig13135:14368-14400(+) | Contig13670:14779-14811(+) | 3 |  |  |
| **Aves** |  |  |  |  |  |
| MediumGroundFinch(geoFor1) | -- | JH739888:20155410-20155442(+) | - | NA |  |
| ZebraFinch(taeGut2) | -- | chr7:17035450-17035482(+) | - | NA |  |
| Budgegerigar(melUnd1) | -- | JH556613:11953860-11953892(-) | - | NA |  |
| Chicken(galGal4) | -- | chr7:15818298-15818330(-) | - | NA | Y |
| Turkey(melGal1) | -- | chr7:16893541-16893573(-) | - | NA |  |
| **Sarcopterygii** |  |  |  |  |  |
| AmericanAlligator(allMis1) | JH739658:224031-224063(-) | JH736379:590770-590802(-) | 3 |  |  |
| PaintedTurtle(chrPic2) | JH584844:1113401-1113433(+) | JH584712:2171341-2171373(+) | 3 |  |  |
| Frog(xenTro3) | GL172862:354043-354075(-) | GL172799:638809-638841(-) | 7 |  | Y |
| Coelacanth(latCha1) | JH126829:1709983-1710015(+) | JH127184:433817-433849(-) | 4 | Y | Y |
| **Holesti Fish** |  |  |  |  |  |
| Spottedgar (lepOcu1) | LG12:15577478-15577509(-) | LG4:15815626-15815646(-) | 3 |  |  |
| **Teleost Fish** |  |  |  |  |  |
| Tetraodon(tetNig2) | -- | chr2:13442915-13442947(-) | - | N | N |
| Fugu(fr3) | -- | chr1:15039680-15039712(-) | - | N | N |
| Medaka(oryLat2) | -- | chr21:24604991-24605023(+) | - | N | N |
| AtlanticCod(gadMor1) | -- | HE566826:374754-374786(-) | - | N | N |
| Zebrafish(danRer7) | -- | chr9:1970782-1970814(-) | - | N | N |
| **Cartilaginous Fish** |  |  |  |  |  |
| ElephantShark(calMal1) | KI636119:205590-205622(-) | KI635868:5106636-5106668(+) | 7 |  |  |
| **Jawless Vertebrates** |  |  |  |  |  |
| Lamprey(petMar2) | -- | -- | - | - | - |
| **Deuterostome** |  |  |  |  |  |
| C.Intestinalis(ci2) | -- | -- | - | - | - |
| **Insect** |  |  |  |  |  |
| D.melanogaster(dm6) | -- | -- | - | - | - |
| **Nematode** |  |  |  |  |  |
| C.elegans(ce6) | -- | -- | - | - | - |

**Supplementary Table 2.** Summary of annotation of HoxC cluster genes in aves and teleosts. Only few HoxC genes are annotated in aves and are located in different contigs. Except chicken, other aves do not have annotated *HoxC11* and *HoxC12* genes. In teleosts, HoxC cluster genes are well annotated and located in the same cluster.

Table Supplemental S2. Summary of annotation of hoxc cluster genes in aves and telesot.
|  | hoxa |  | hoxb |  | hoxc |  | hoxd |  | Annotation |  | Same | Annotation |  | Same |
| --- | --- | --- | --- | --- | --- | --- | --- | --- | --- | --- | --- | --- | --- | --- |
|  | Genes | Chr | Genes | Chr | Genes | Chr | Genes | Chr | hoxc11 | hoxc12 | Chr | hoxd11 | hoxd12 | Chr |
| Aves |  |  |  |  |  |  |  |  |  |  |  |  |  |  |
| ZebraFinch | 9 | 1 | 5 | 1 | 2 | 2 | 5 | 2 | NO | NO | NA | YES | YES | YES |
| Chicken | 8 | 1 | 9 | 1 | 5 | 5 | 7 | 1 | YES | YES | Diff | YES | YES | YES |
| Turkey | 4 | 1 | 4 | 2 | 2 | 1 | 5 | 1 | NO | NO | NA | YES | YES | YES |
| Teleost |  |  |  |  |  |  |  |  |  |  |  |  |  |  |
| Tetraodon | 13 | 2 | 9 | 2 | 11 | 2 | 6 | 1 | YES | YES | YES | YES | YES | YES |
| Fugu | 9 | 3 | 7 | 3 | 6 | 1 | 4 | 1 | NO | YES | NA | NO | YES | NA |
| Atlantic Cod | 12 | 5 | 9 | 3 | 10 | 1 | 6 | 1 | YES | YES | YES | YES | YES | YES |
| Zebrafish | 12 | 2 | 15 | 2 | 15 | 2 | 7 | 1 | YES | YES | YES | YES | YES | YES |

**Supplementary Table 3.**
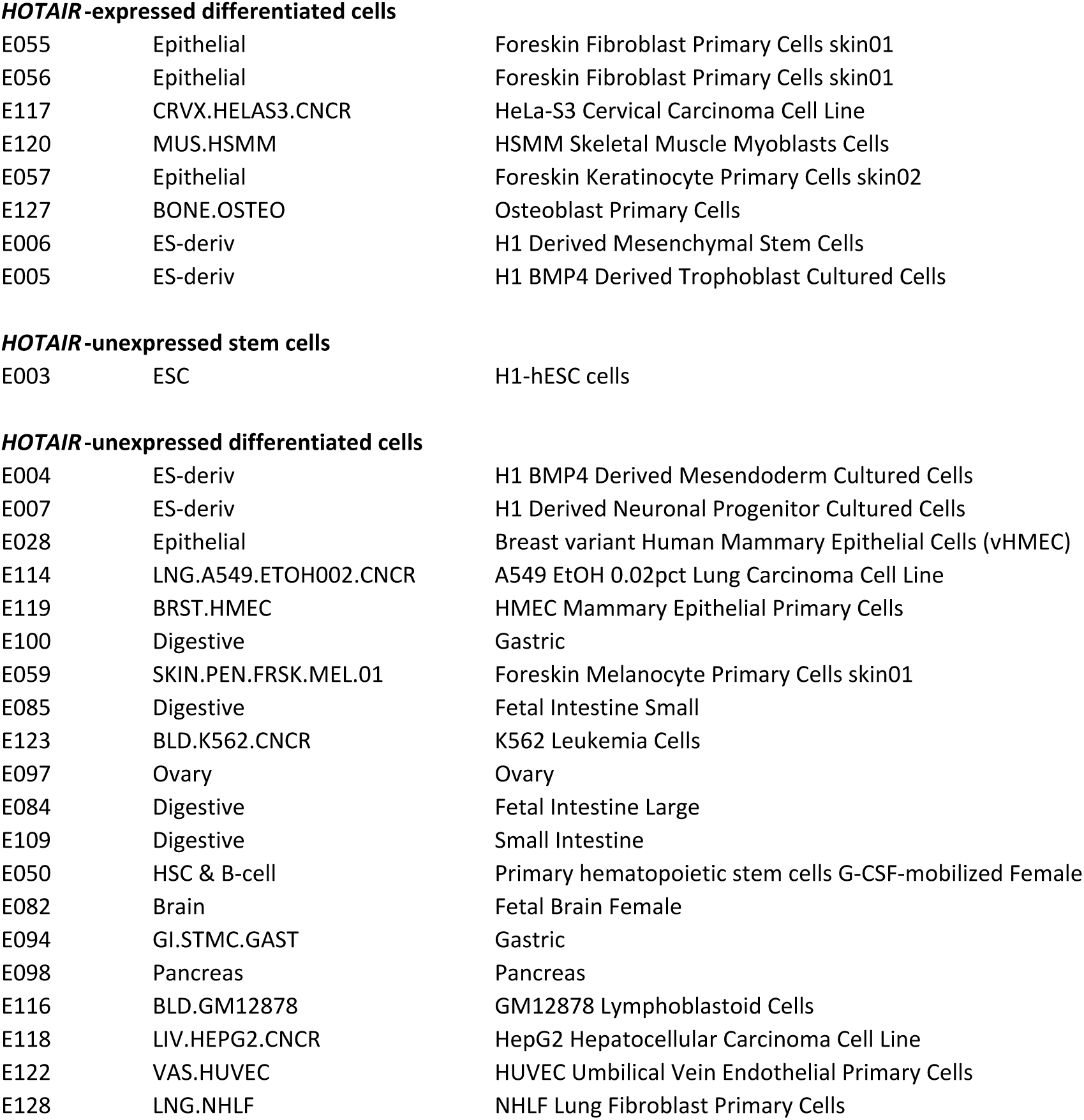
List of cell lines from Roadmap Epigenome samples that are classified into three groups based on *HOTAIR* expression from RNA-seq.

**HOTAIR-expressed differentiated cells**
|  |  |  |
| --- | --- | --- |
| E055 | Epithelial | Foreskin Fibroblast Primary Cells skin01 |
| E056 | Epithelial | Foreskin Fibroblast Primary Cells skin01 |
| E117 | CRVX.HELAS3.CNCR | HeLa-S3 Cervical Carcinoma Cell Line |
| E120 | MUS.HSMM | HSMM Skeletal Muscle Myoblasts Cells |
| E057 | Epithelial | Foreskin Keratinocyte Primary Cells skin02 |
| E127 | BONE.OSTEO | Osteoblast Primary Cells |
| E006 | ES-deriv | H1 Derived Mesenchymal Stem Cells |
| E005 | ES-deriv | H1 BMP4 Derived Trophoblast Cultured Cells |
| **HOTAIR-unexpressed stem cells** |  |  |
| E003 | ESC | H1-hESC cells |
| **HOTAIR-unexpressed differentiated cells** |  |  |
| E004 | ES-deriv | H1 BMP4 Derived Mesendoderm Cultured Cells |
| E007 | ES-deriv | H1 Derived Neuronal Progenitor Cultured Cells |
| E028 | Epithelial | Breast variant Human Mammary Epithelial Cells (vHMEC) |
| E114 | LNG.A549.ETOH002.CNCR | A549 EtOH 0.02pct Lung Carcinoma Cell Line |
| E119 | BRST.HMEC | HMEC Mammary Epithelial Primary Cells |
| E100 | Digestive | Gastric |
| E059 | SKIN.PEN.FRSK.MEL.01 | Foreskin Melanocyte Primary Cells skin01 |
| E085 | Digestive | Fetal Intestine Small |
| E123 | BLD.K562.CNCR | K562 Leukemia Cells |
| E097 | Ovary | Ovary |
| E084 | Digestive | Fetal Intestine Large |
| E109 | Digestive | Small Intestine |
| E050 | HSC & B-cell | Primary hematopoietic stem cells G-CSF-mobilized Female |
| E082 | Brain | Fetal Brain Female |
| E094 | GI.STMC.GAST | Gastric |
| E098 | Pancreas | Pancreas |
| E116 | BLD.GM12878 | GM12878 Lymphoblastoid Cells |
| E118 | LIV.HEPG2.CNCR | HepG2 Hepatocellular Carcinoma Cell Line |
| E122 | VAS.HUVEC | HUVEC Umbilical Vein Endothelial Primary Cells |
| E128 | LNG.NHLF | NHLF Lung Fibroblast Primary Cells |

**Supplementary Table 4.** Enrichment of transcription factors bound on the *HOTAIR* CNE and promoter.

**Table Supplemental S4: Enrichment of transcription factors (TFs) bound on *HOTAIR* CNE and promoter.**
| chr | start | end | region | - | strand | chr | start_TFPeak | End_TFPeak | TF | PeakScore | Celllines |
| --- | --- | --- | --- | --- | --- | --- | --- | --- | --- | --- | --- |
| chr12 | 54359507 | 54360239 | HotairCNE | 1000 | - | chr12 | 54359825 | 54360441 | CHD1 | 261 | H1-hESC |
| chr12 | 54359507 | 54360239 | HotairCNE | 1000 | - | chr12 | 54359540 | 54359800 | TCF12 | 178 | H1-hESC |
| chr12 | 54359507 | 54360239 | HotairCNE | 1000 | - | chr12 | 54359918 | 54360194 | EGR1 | 319 | H1-hESC |
| chr12 | 54359507 | 54360239 | HotairCNE | 1000 | - | chr12 | 54359520 | 54359844 | SP1 | 240 | H1-hESC |
| chr12 | 54359507 | 54360239 | HotairCNE | 1000 | - | chr12 | 54359673 | 54359803 | SRF | 631 | H1-hESC |
| chr12 | 54359507 | 54360239 | HotairCNE | 1000 | - | chr12 | 54359955 | 54360245 | ZNF143 | 158 | H1-hESC |
| chr12 | 54359507 | 54360239 | HotairCNE | 1000 | - | chr12 | 54359398 | 54360436 | CTBP2 | 262 | H1-hESC |
| chr12 | 54359507 | 54360239 | HotairCNE | 1000 | - | chr12 | 54360179 | 54360920 | SUZ12 | 224 | H1-hESC,NT2-D1 |
| chr12 | 54359507 | 54360239 | HotairCNE | 1000 | - | chr12 | 54359205 | 54365366 | EZH2 | 1000 | GM12878,H1-hESC,HepG2,HUVEC,NH-A,NHLF |
| chr12 | 54362255 | 54362956 | HotairPromoter | 1000 | - | chr12 | 54362190 | 54362660 | POLR2A | 204 | PANC-1,SK-N-MC |
| chr12 | 54362255 | 54362956 | HotairPromoter | 1000 | - | chr12 | 54362520 | 54363116 | TCF7L2 | 426 | PANC-1 |
| chr12 | 54362255 | 54362956 | HotairPromoter | 1000 | - | chr12 | 54359205 | 54365366 | EZH2 | 1000 | GM12878,H1-hESC,HepG2,HUVEC,NH-A,NHLF |
| chr12 | 54362255 | 54362956 | HotairPromoter | 1000 | - | chr12 | 54362041 | 54362337 | GATA3 | 234 | T-47D |
| chr12 | 54362255 | 54362956 | HotairPromoter | 1000 | - | chr12 | 54361277 | 54362537 | SUZ12 | 167 | H1-hESC,NT2-D1 |
| chr2 | 176968657 | 176969389 | HoxD11CNE | 1000 | + | chr2 | 176969091 | 176969421 | CTCF | 526 | A549,Fibrobl,Gliobla,HEK293,SK-N-SH_RA |
| chr2 | 176968657 | 176969389 | HoxD11CNE | 1000 | + | chr2 | 176968367 | 176975129 | EZH2 | 1000 | H1-hESC,HepG2,HMEC,HSMM,HSMMtube,HUVEC,NH-A,NHDF-Ad,NHEK,NHLF |
| chr2 | 176968657 | 176969389 | HoxD11CNE | 1000 | + | chr2 | 176968464 | 176968800 | POLR2A | 382 | SK-N-MC |
| chr2 | 176968657 | 176969389 | HoxD11CNE | 1000 | + | chr2 | 176968988 | 176969498 | POLR2A | 471 | A549,SK-N-MC,Gliobla |
| chr2 | 176968657 | 176969389 | HoxD11CNE | 1000 | + | chr2 | 176968889 | 176969429 | EP300 | 381 | A549 |
| chr2 | 176968657 | 176969389 | HoxD11CNE | 1000 | + | chr2 | 176969019 | 176969197 | FOSL2 | 813 | A549 |
| chr2 | 176968657 | 176969389 | HoxD11CNE | 1000 | + | chr2 | 176968813 | 176969389 | TCF12 | 356 | A549 |
| chr2 | 176968657 | 176969389 | HoxD11CNE | 1000 | + | chr2 | 176969043 | 176969353 | SUZ12 | 96 | H1-hESC |
| chr2 | 176968657 | 176969389 | HoxD11CNE | 1000 | + | chr2 | 176968819 | 176969335 | TCF7L2 | 185 | HEK293 |
| chr2 | 176968657 | 176969389 | HoxD11CNE | 1000 | + | chr2 | 176968726 | 176969412 | KAP1 | 329 | HEK293,U2OS |

**Supplementary Table 5.**
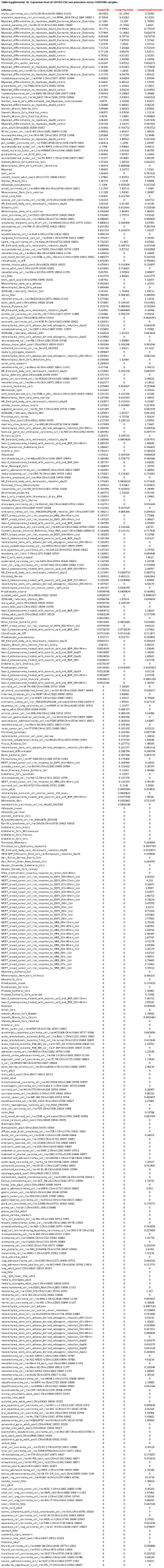
Expression level of the *HOTAIR* CNE and promoter across FANTOM5 samples.

---

**Supplementary Figure 1.**
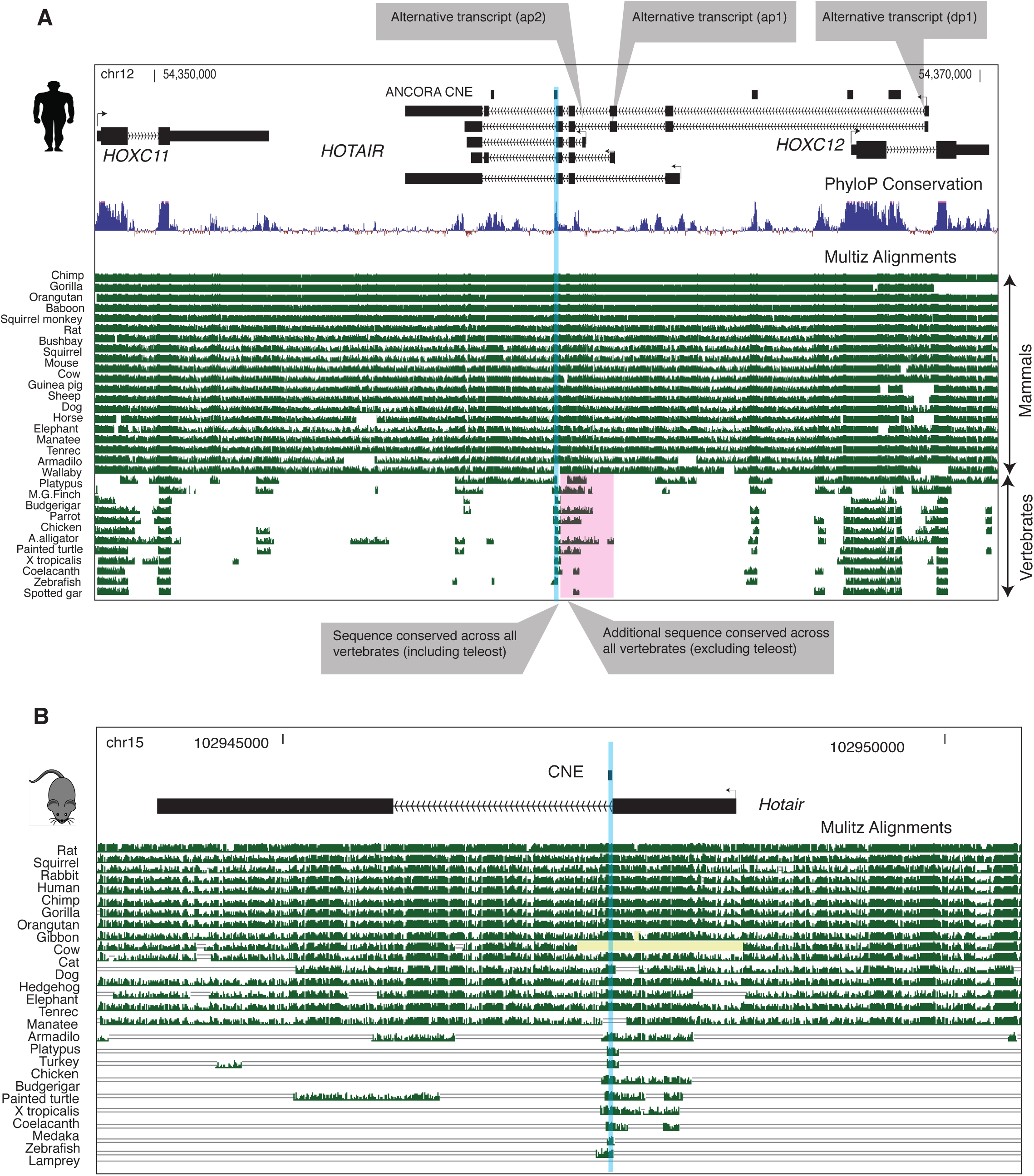
A genome browser view of *HOTAIR* in human and mouse to show sequence conservation. (**A**) Conserved noncoding elements (CNEs) track from ANCORA browser is shown as horizontal black bars on top. CNE that is conserved across all mammals and vertebrates is highlighted. CNE is located eight nucleotides away from splice site. CNE is mapped across all vertebrates, but flanking sequences (corresponding *HOTAIR* exons) are mapped in other fish and tetrapod (excluding teleost). (**B**) Structure of mouse *Hotair* transcript is different than in human. Homolog of CNE is also located eight nucleotides away from splice site of *Hotair* transcript. CNE is conserved across mammals and vertebrates.

**Supplementary Figure 2.**
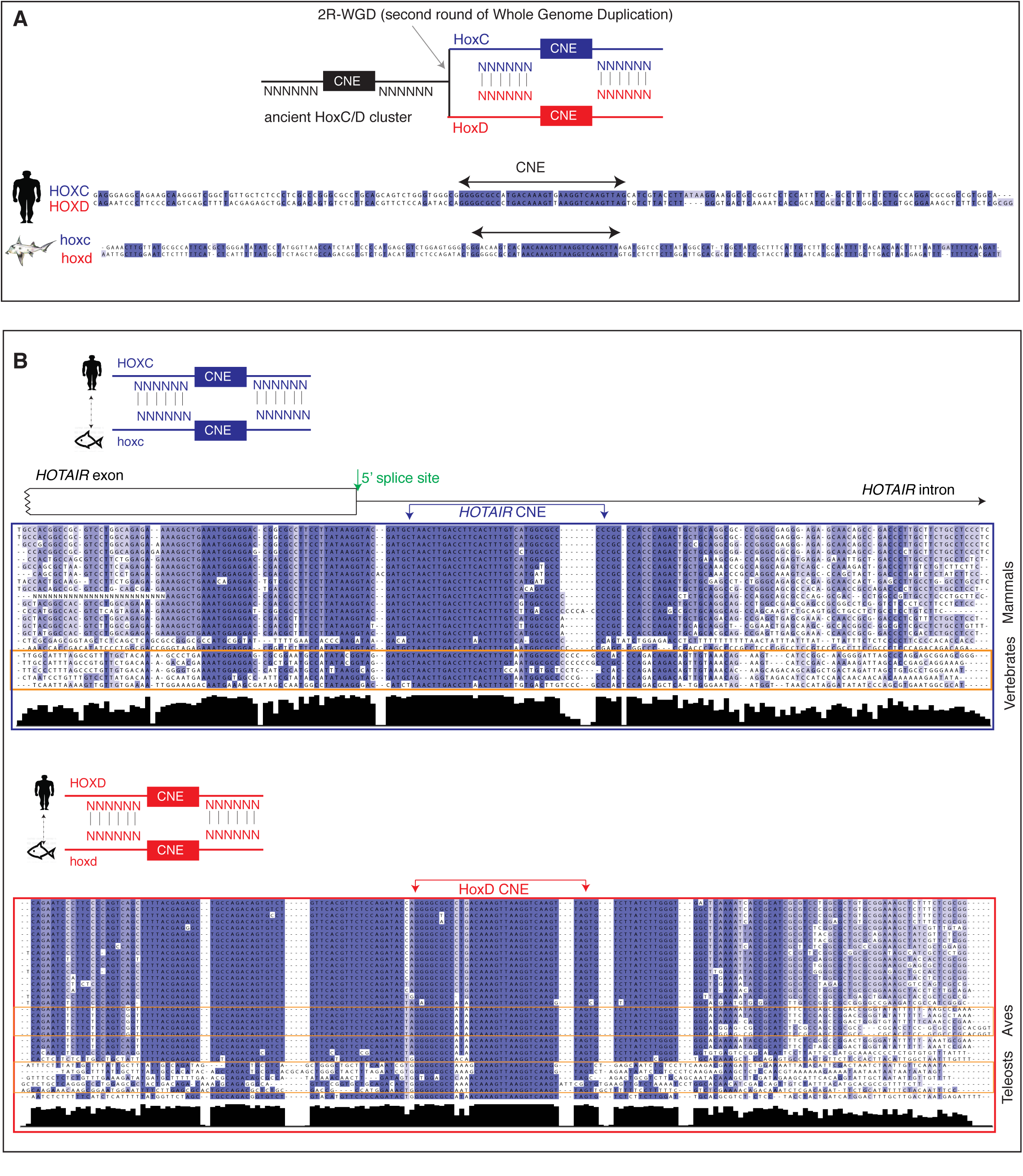
Alignment of CNEs and flanking sequences across HoxC and HoxD clusters within and across species. (**A**) Schematic representation on the origin of HoxC and HoxD cluster from the ancestral HoxC/D cluster after second round of whole genome duplication (2R-WGD). Alignment of flanking sequences around paralogous CNEs, in human and elephant shark, show little conservation despite both sequences are duplicated from the same ancestral sequences. (**B**) Schematic representation of alignment HoxC and HoxD CNE and flanking sequences across vertebrates. Alignment of 70 nucleotides flanking the *HOTAIR* CNE (across 22 vertebrates (excluding aves and teleosts)) and HOXD CNE (across 32 vertebrates) show conservation of flanking sequences across vertebrates. Species are aligned in the same order as in Supplementary Figure S1A.

**Supplementary Figure 3.**
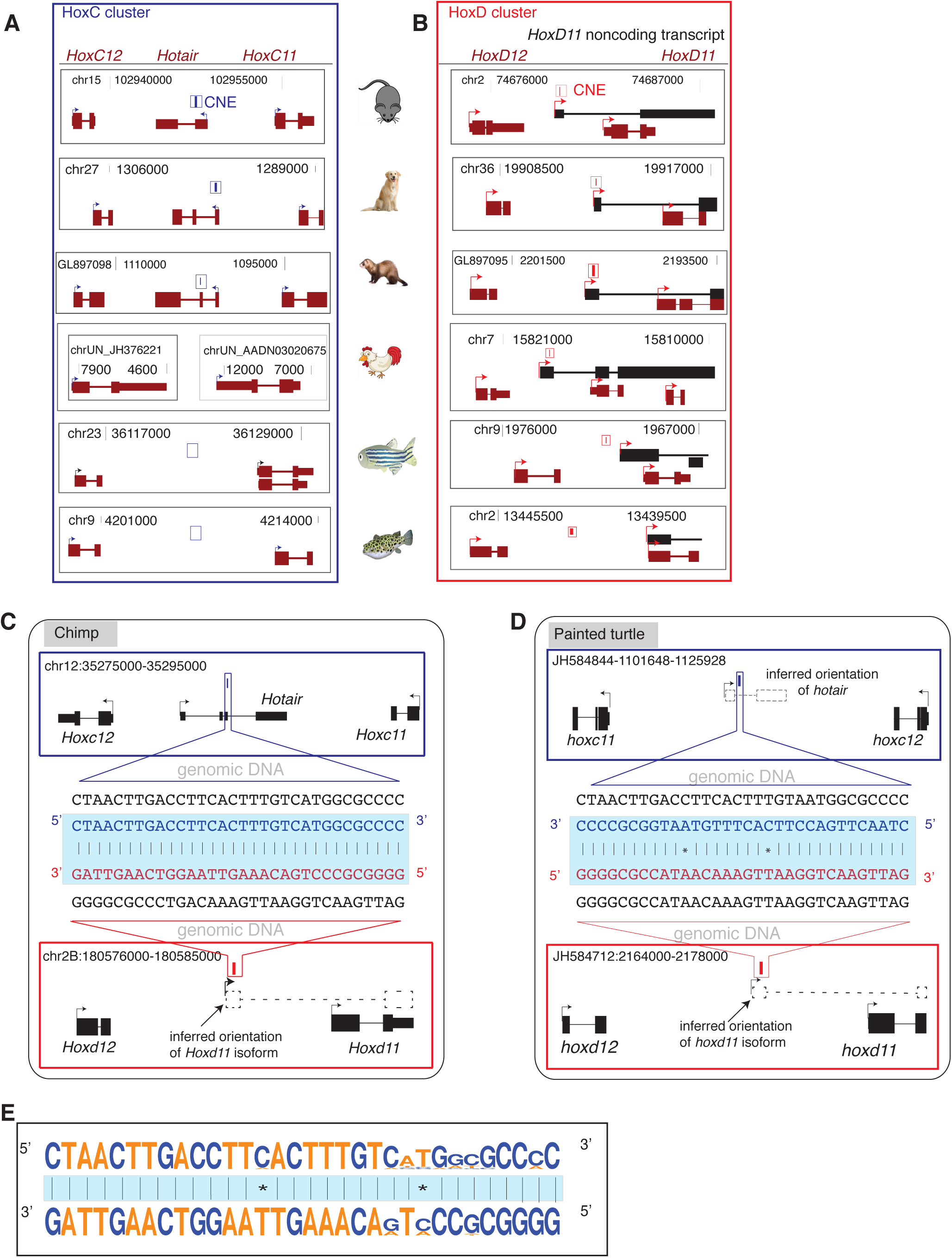
Evidence of *HOTAIR* and *HoxD11* noncoding transcript across multiple species and sequence complementarity of paralogous CNEs. (**A**) The *HOTAIR* CNE is represented by rectangular blue bar. A homolog of the *HOTAIR* CNE is undetected in chicken. Intergenic region between *hoxc11* and *hoxc12* remains unassembled as *hoxc11* and *hoxc12* genes are assembled in different contigs. Zebrafish *hoxc11* and *hoxc12* is assembled but lacks the CNE. (**B**) The HoxD CNE is represented by rectangular red bar. *HoxD11* noncoding transcript is detected across tetrapods but not in teleosts. (**C-D**) Schematic representation to show how the orientation of missing *Hotair* and *HoxD11* noncoding transcripts were inferred. *HOTAIR* is antisense to *HoxC11* genes across species, and thus the same convention was used to infer the orientation of *Hotair* CNE. HoxD11 noncoding transcript is an alternative splice variant of *HoxD11* across multiple species; thus, the same convention was used. Expected transcripts are represented by dashed rectangular boxes and lines and arrows indicate transcript orientation. The CNE sequences are zoomed in on to show the genomic DNA. Aligning sequences in 5’ to 3’ orientation show paralogous CNEs are complementary. (**E**) Sequence logos of *HOTAIR* CNE and HoxD CNE show paralogous CNEs exhibit sequence complementarity in transcribed orientation.

**Supplementary Figure 4.**
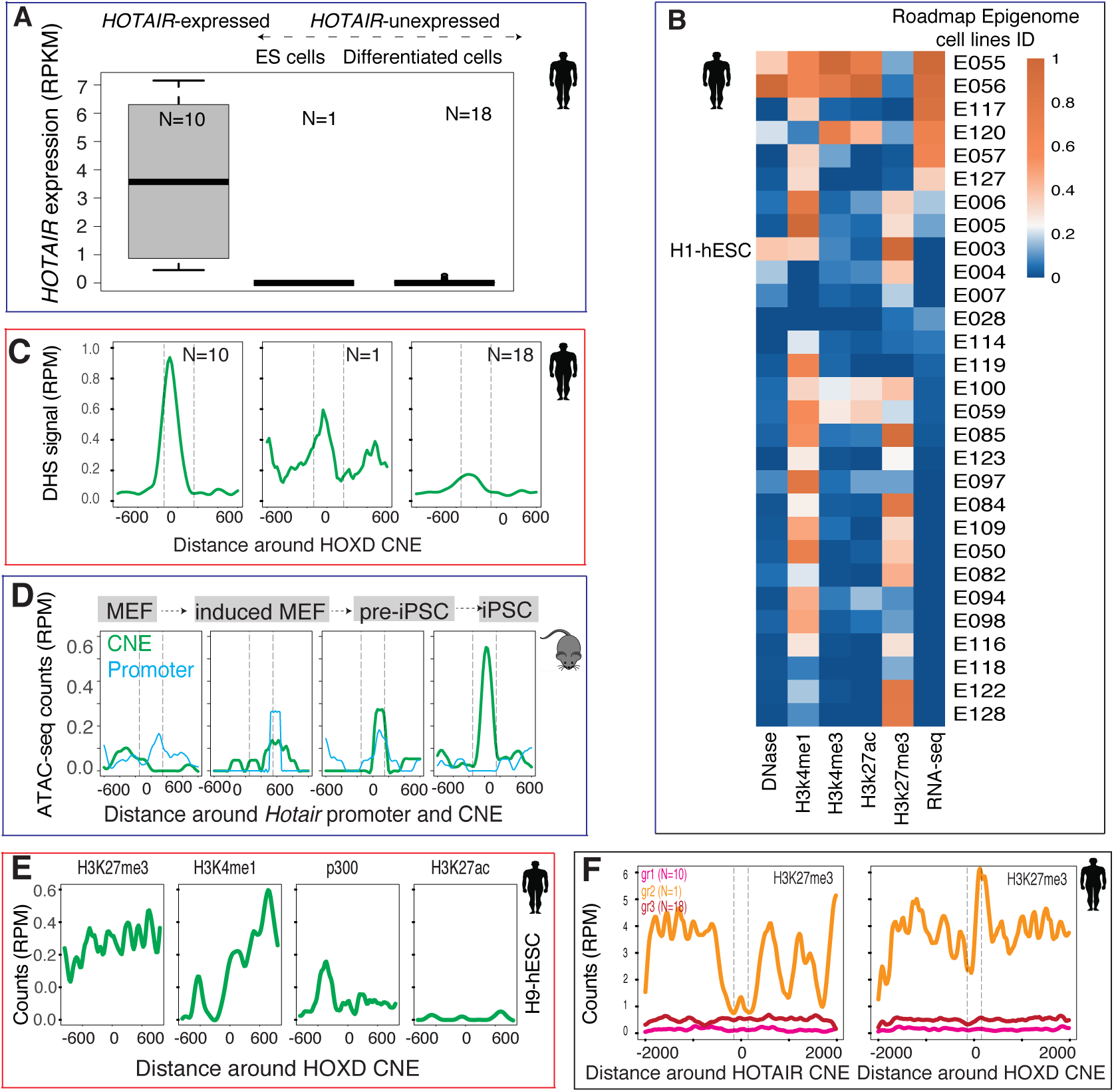
Chromatin environment of the *HOTAIR* CNE. (**A**) Twenty-nine cell lines were separated into *HOTAIR*-expressed and *HOTAIR*-unexpressed cell lines using a threshold of 0.5 RPKM (reads per kilobase per million mapped reads). *HOTAIR*-unexpressed cell lines were further separated into stem cells (N=1) and terminally differentiated cells (N=18). (**B**) Heatmaps show the normalized read counts in 250 nucleotides flanking *HOTAIR* CNE across 29 cell lines from Roadmap Epigenome Consortium. Cell line IDs are the same as provided by Roadmap Epigenome Consortium. (**C**) DHS signals around the HOXD CNE across three groups (*HOTAIR*-expressed cells, *HOTAIR*-unexpressed stem cells and *HOTAIR*-unexpressed cells) show similar trends as that of the *HOTAIR* CNE. Y-axis represents normalized read counts in reads per million (RPM). (**D**) Open chromatin state of mouse *Hotair* CNE is dynamically acquired during reprogramming of mouse embryonic fibroblast to iPSC. (**E**) Human HOXD CNE is not a poised enhancer as it lacks p300, H3K4me1 and bimodal H3K27me3 peaks in H9-hESC cell line. (**F**) Pattern of H3K27me3 marks around *HOTAIR* CNE across three groups are different. H3K27me3 patterns around the HOXD CNE show similar enrichment across three groups.

**Supplementary Figure 5.**
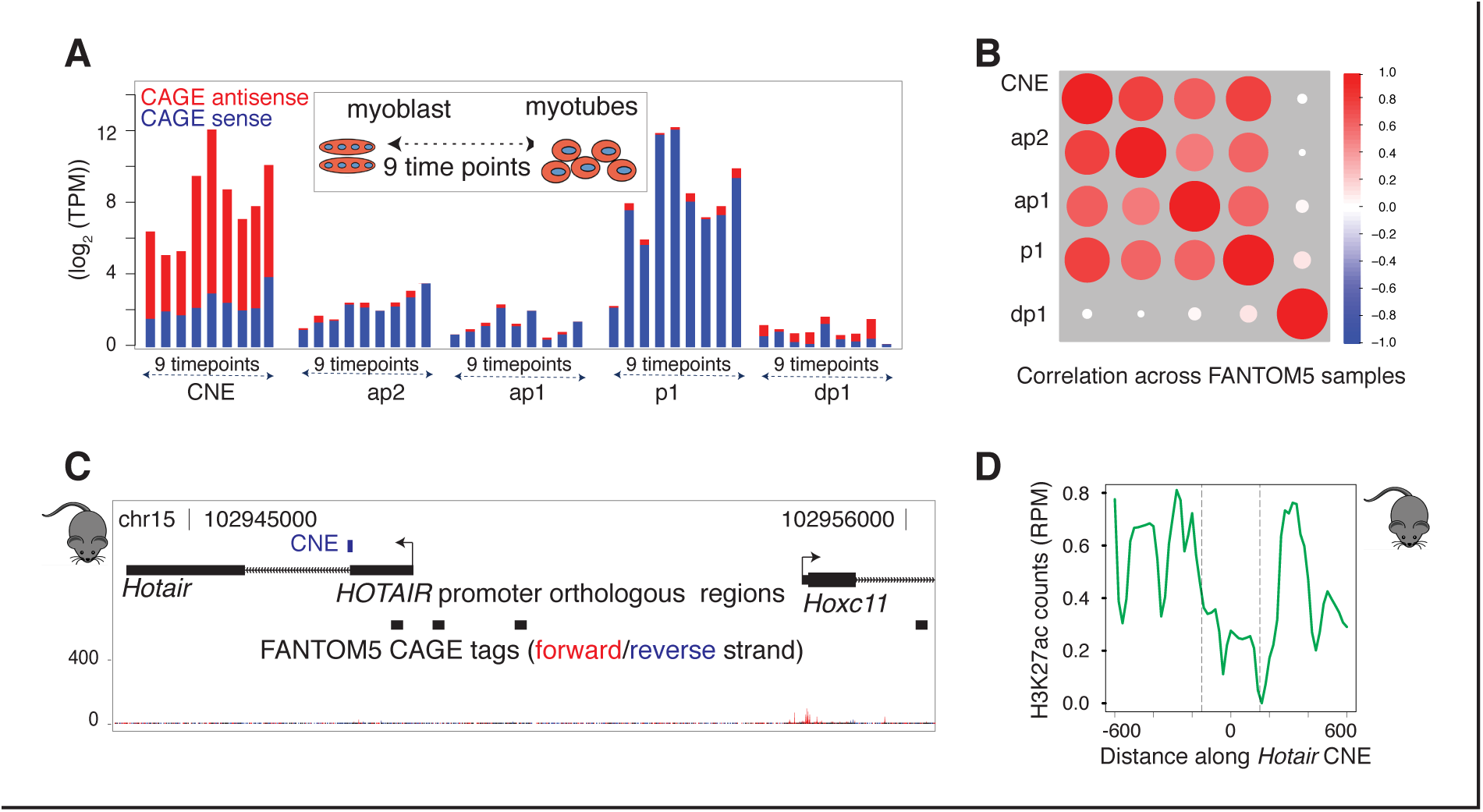
Transcriptional dynamics of *HOTAIR* promoters. (**A**) Expression levels of *HOTAIR* promoter, alternative promoters and CNE across nine different stages during differentiation from myoblast to myotube. (**B**) Expression correlation of *HOTAIR* promoter, alternative promoters and CNE across FANTOM5 samples are positively correlated except for distal promoter (dp1). (**C**) No significant CAGE tags are identified on mouse *Hotair* across FANTOM5 samples as it lacks tissues (embryonic hindlimbs, genital tubercle and a piece of trunk corresponding to sacro-caudal region) where Hotair is expressed. Orthologs of promoter region of HOTAIR are aligned to mouse. (**D**) *Hotair* CNE is flanked by bidirectional H3K27ac on mouse hindlimbs (E10.5 days).

**Supplementary Figure 6.**
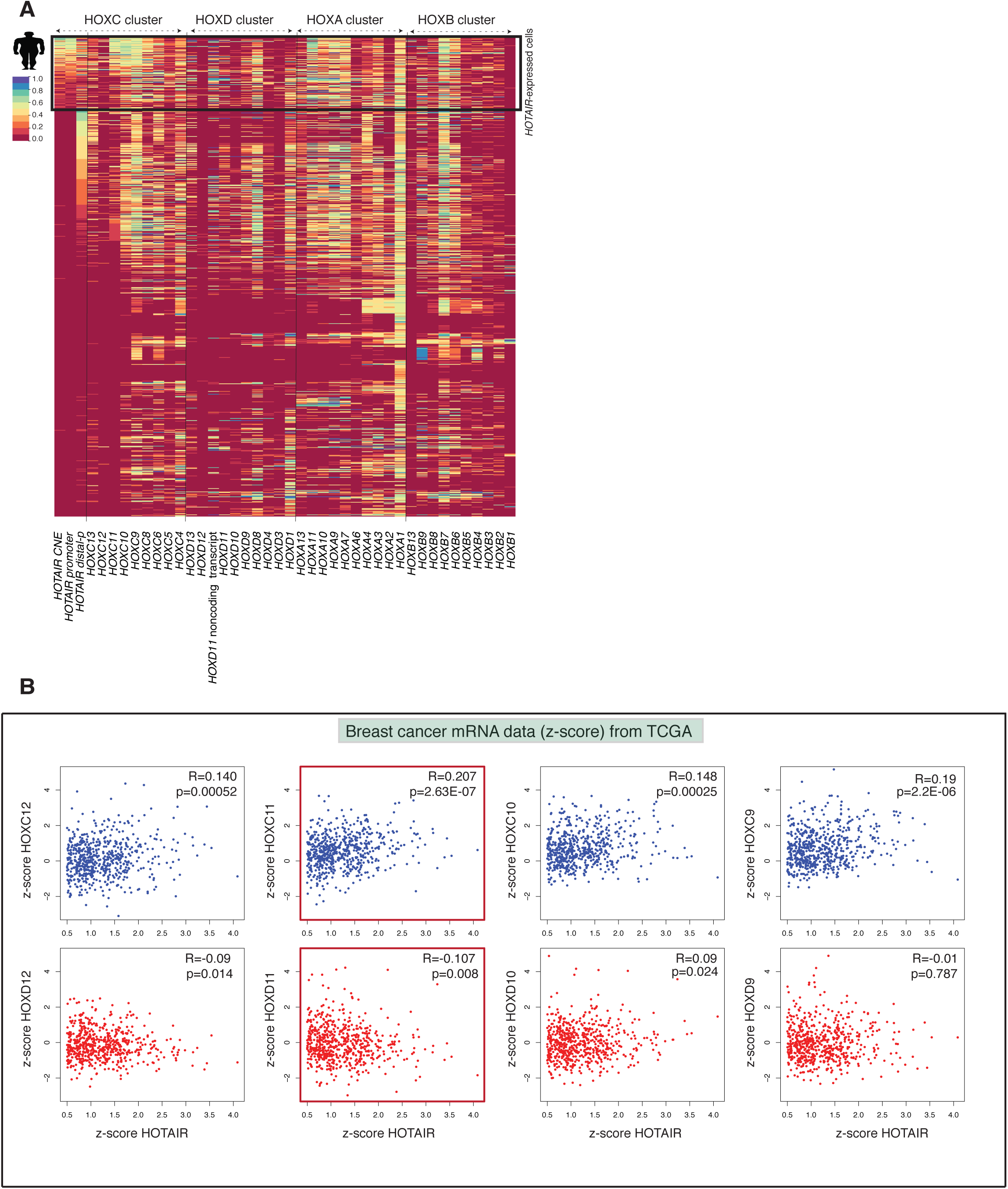
Co-expression analysis of *HOTAIR* with four HOXC clusters genes across FANTOM5 samples. (**A**) Heatmap shows the expression level of *HOTAIR* and four HOX cluster genes. Cell types are sorted based on *HOTAIR* CNE and *HOTAIR* distal promoter expression. Expression levels are scaled between 0-1 across each column, where the most highly expressed gene is assigned 1. (**B**) Expression levels of *HOTAIR* with HOXC and HOXD cluster posterior genes on individual breast cancer patients show a trend of positive correlation with HOXC cluster genes and a trend of weak negative correlation with HOXD cluster genes.

